# Open filament tracks drive Bacteroidota gliding motility

**DOI:** 10.64898/2026.11.02.763665

**Authors:** Helen L. Miller, Rachel L. Jones, Andreas Kjӕr, Simone Coppola, Ben C. Berks, Richard M. Berry

## Abstract

Bacteria of the phylum Bacteroidota glide across surfaces using adhesins that attach to the substratum and are propelled along the cell body by an underlying track. The track is currently thought to form a single closed loop around the cell. Here we use 3D super-resolution tracking of single adhesin molecules in *Flavobacterium johnsoniae* to characterize adhesin movements in unprecedented detail. Our data overturn multiple aspects of the single closed loop model by showing that adhesins follow non-repeating trajectories that cross, make sharp turns, and exhibit reversals and circular paths. We develop a new model for Bacteroidota gliding in which the track network comprises multiple independently moving, open filaments. This supramolecular organization resembles that of other moving bacterial filaments such as MreB and FtsZ, suggesting that open filaments are a common organising principle in the bacterial cytoskeleton.

## Introduction

Gliding is a form of surface-associated bacterial motility in which the organism attaches to a substratum through cell surface adhesin proteins that are actively propelled along the body of the cell^1,2^. The most widely distributed gliding system is found in bacteria of the Gram-negative phylum Bacteroidota (formerly Bacteroidetes). Gliding is the only type of motility utilized by this major phylum and most free-living Bacteroidota possess the genes for the gliding apparatus^3-5^. Gliding Bacteroidota are found in association with many solid surfaces including decaying plant fibres, soil particles, plant roots, the surface of the oral cavity, and marine snow, providing a rationale for their capacity for surface-linked motility^6^. Their gliding motility is implicated in the suppression of plant disease by root-associated bacteria^7^, in the virulence of pathogens of farmed fish^8^, in predatory behaviour^9,10^, in the dissemination of non-motile bacteria and phages through biofilms^11,12^, and in collective cell motion^13,14^ that can lead to colony structuring with potential biotechnological applications^15^. Gliding is a property of the *Capnocytophaga* species found in the oral cavity, which include both commensals and opportunistic pathogens^11^.

Despite the importance of gliding motility to the Bacteroidota, the mechanism of gliding is still poorly understood. Bacteroidota gliding motility has been most intensively studied using *Flavobacterium johnsoniae* as a model system^4^. *F. johnsoniae* moves at speeds up to 2 μm s^-1^ on hard surfaces overlaid with aqueous media. This movement is mediated by cell surface adhesins that flow in helical paths back and forth along the cell body^16,17^. Because the adhesins are moving relative to the cell body, immobilization of the adhesins by attachment to the substratum propels the cells forward in a corkscrewing motion along the long axis of the cell. This corkscrewing motion has been directly visualized by imaging bent filamented cells^17^ or by using stationary outer membrane proteins as fiducial markers^18,19^. Because the cell must be able to achieve forward motion even though the adhesins are moving in both directions along the cell body, it is proposed that only those adhesins moving in the rearward direction are competent to bind to the gliding substratum^16,20^ (Fig. 1a). The energy for moving the adhesins comes from the protonmotive force (PMF) across the inner membrane of the cell^16,21,22^.

**Figure 1.**
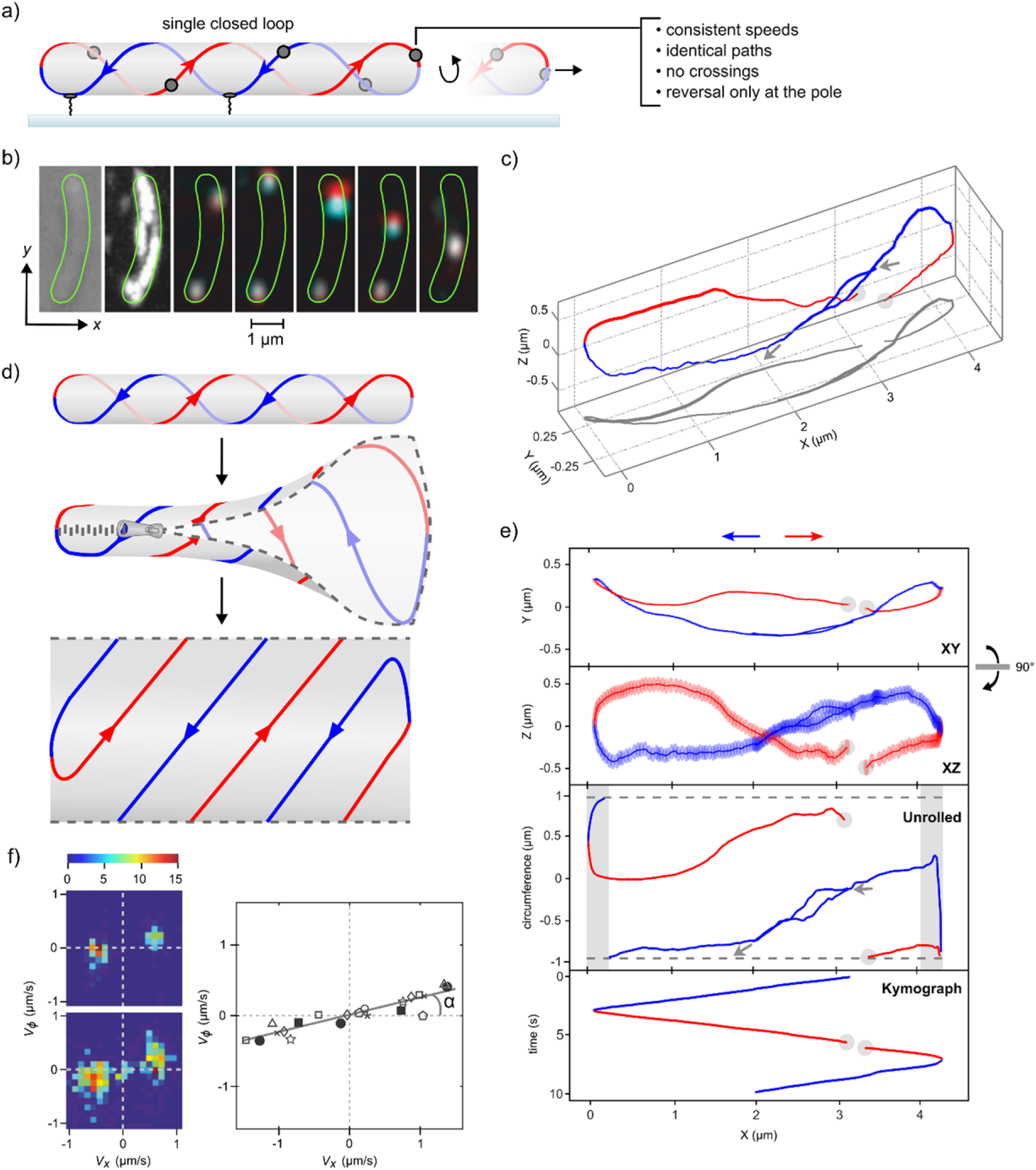
Analysis of the trajectories of single SprB adhesin molecules. To aid visualization, rightwards and leftwards movements are coloured red and blue respectively. (a) Adhesin paths and their expected properties in the single closed loop model. Rearward-moving adhesins bind the substratum to propel gliding cells. (b) A fluorophore-labelled SprB adhesin molecule moving in an immobilized cell illustrating how depth is encoded as differential movement in the parallax-shifted y axis between two parallax views of the cell. Left to right: brightfield, fluorescence maximum-projection of one view, overlaid parallax views (red, cyan) at ^1^/6 s intervals. An indicative cell outline is shown in green. A second stationary molecule at the bottom of the image photobleaches before the last frame. See also Video S1. (c) 3D plot of the adhesin trajectory derived from the cell in **b**. The start and end points (grey arrows) and a gap where localization failed (grey circle) are indicated. Line thickness increases with distance (*z*) from the coverslip. The projection of the trajectory onto the coverslip is shown in grey. (d) Schematic showing the relation between the 3D data in **c** and the corresponding Unrolled plot in **e**. (e) The trajectory in **c** projected onto the XY (coverslip) or XZ planes, in unrolled cylindrical co-ordinates, and the cell long axis position versus time (“Kymograph”). XY and XZ panels show measurement error as translucent ellipses. Start and end of the trajectory shown as grey arrows in the Unrolled plot. Grey overlay indicates regions of the cell where the cylindrical projection is a distorted representation of the spherical end cap. (f) Left: Velocity density distributions for the cell in **b,c,e** (upper) and for another cell (lower). Colour bar indicates numbers of observations. Right: Peaks in velocity distributions from 8 cells (one symbol per cell) with linear fit (grey line). Velocity components (VX, Vφ) defined as the per-frame change in cylindrical co-ordinates divided by the frame time. α = 13.7 ± 1.8° represents the average angle of adhesin movement. See Fig. S1 for the expected appearance of velocity distributions for different adhesin behaviours.

SprB is the major gliding adhesin of *F. johnsoniae*^23^. SprB is a ∼150 nm long filamentous protein^16^ that is anchored to the cell surface by the outer membrane-spanning protein SprF^24^. SprF is in turn connected through a disulfide linkage to a moving track structure located at the periplasmic face of the outer membrane^25^. A motor complex in the inner membrane transduces the energy of the PMF into mechanical energy and transmits this across the periplasm to move the gliding track at the outer membrane^25-28^.

Our current understanding of the organization of the gliding track network in *F. johnsoniae* has been inferred from experiments in which the movements of adhesin molecules were studied after labelling them with polystyrene or gold beads, or with fluorescent antibodies^16,17,23,29,30^. These experiments have led to a model in which the adhesins move at constant velocity along a single helical track which forms a closed loop that runs the full length of the cell once in either direction (Fig. 1a)^16,20,31,32^. However, more complex models have also been envisaged in which multiple closed tracks are present that would need to cross each other at the cell ends^21^ or multiple tracks moving in either direction are bundled together^19^.

To resolve the organization of the gliding tracks and other questions about the mechanism of gliding motility we have used 3-dimensional (3D) tracking of single *F. johnsoniae* SprB adhesin molecules to determine the trajectories of moving adhesins in unprecedented detail. We have been able to observe single adhesins making multiple circuits up and down the cell and for the first time resolve the paths of adhesin molecules at the cell ends. Our results show that adhesins do not normally follow the same path on successive circuits of the cell and that tracks follow multiple paths at the cell ends, including sharp turns and paths that cross each other. We further show that adhesin movements can deviate markedly from the idealized behaviour assumed in current models, including different speeds within the same cell, paths that do not reach the cell end, and paths that are circular rather than helical. In light of these findings, we present a revised model for the organization of the gliding track network in which there are multiple open-filament tracks.

## Results

### Experimental strategy

To achieve a step change in our understanding of the *F. johnsoniae* gliding track network, we implemented long-duration, high-precision 3D tracking of individual adhesin molecules moving along an experimentally immobilized cell body.

We labelled individual SprB molecules with the bright and highly photostable fluorophore Janelia Fluor 646 (JF646) using a HaloTag domain fused to the N-terminus of the adhesin^28^. Direct labelling of the adhesin in this way ensures that we are tracking single adhesin molecules and contrasts with earlier studies^16,17,23,29,30^ in which the adhesins were labelled with polyvalent ligands (beads or antibodies) that are likely to bind simultaneously to multiple adhesin molecules. The covalent linkage of the fluorophore to the adhesin in our experiments also avoids the possibility that the label could move between adhesins during imaging. We employed sparse labelling of SprB to allow individual adhesin molecules to be unambiguously resolved and tracked. Because individual adhesins were imaged, their trajectories could be determined with high precision using single molecule localization (‘super-resolution’ accuracy). The high photostability of JF646 allowed the adhesins to be tracked for periods of 10 s or more, sometimes permitting single adhesins to be followed through more than one circuit of the cell. We imaged cells immobilized between agarose and glass, allowing adhesin movements to trace out the positions of the tracks in the reference frame of a static cell.

We followed the movements of the adhesin molecules in 3D. This enabled us to define the trajectory of the adhesins without ambiguity, and for the first time allowed us to determine the adhesin paths at the ends of the cell. 3D tracking was carried out using parallax imaging^33^ implemented on a custom-built microscope. The parallax technique, which is equivalent to stereoscopic vision, recovers the depth of a particle (the *z* co-ordinate) from differential shifts along one axis (assigned here as the *y* co-ordinate) in two views of the particle obtained via opposite halves of the objective back focal plane (see Methods for more details). This technique is suitable for the 3D imaging of moving particles because determining the *z*-localization does not involve fitting the shape of the fluorescence envelope and is thus insensitive to motion blur of the point spread function. The resulting localisations were defined as (*x, y, z*) points in the microscope coordinate frame.

### 3D trajectories of SprB molecules follow left-handed helical paths

Figure 1b-e shows an example of an adhesin trajectory obtained using our experimental system. In this particular example two fluorescent adhesin molecules were observed in the same cell (Fig. 1b). One remained stationary, the other made one complete circuit of the cell and started a second circuit approximately 1 s before the trajectory was terminated by photobleaching of the fluorophore. We converted he raw data from the moving adhesin in Fig. 1b into a ‘super-resolution’ 3D trajectory (Fig. 1c, see Videos S1, S2 for an illustration of this process) with single molecule localisations plotted as 7-point running averages (for a more detailed description see Methods). The initial (*x, y, z*) coordinates were transformed to give (X, Y, Z) coordinates (see Methods) such that the X axis is defined as the long axis of the cell. To enable comparison with previous fluorescence studies of adhesin movements^16,19^ we show the corresponding projection onto the plane of the microscope coverslip (Fig. 1e, “XY”), reducing the data to 2 dimensions. We also show a 2D projection perpendicular to this XY projection (Fig. 1e, “XZ”) in which the cell is viewed parallel to the coverslip. Both the “XY” and “XZ” projections have the measurement error indicated as translucent ellipses at each point. Note that the measurement error is largest in the *z* direction (which is equivalent to Z).

To facilitate the inspection and analysis of the 3D adhesin trajectories we take advantage of the fact that cells are approximately cylindrical and transform the 3D data into a cylindrical co-ordinate plot (Fig. 1e, “Unrolled”). This process is equivalent to unzipping the cell surface parallel to the long axis and laying it flat (Fig. 1d). When presenting trajectories this way we locate the line of unzipping wherever best illustrates chosen features of the data. To further aid trajectory analysis it is useful to also plot the corresponding kymograph showing how the position of the adhesin along the long axis of the cell changes with time (Fig. 1e, “Kymograph”). The datasets used in the remainder of this work were curated to remove short, poorly tracked, or stationary trajectories, retaining only long mobile trajectories for analysis and display (see Methods for more details). Note that it is not possible to track adhesins unambiguously when two or more coincide in 2D, nor when the signal-to-noise ratio dips below the detection threshold. Our analysis assigns different trajectories either side of such gaps (e.g. Fig 1c,e, grey circles), but where we are confident that both trajectories arise from the same adhesin we manually assign the trajectories to a single adhesin and plot them as one.

We plotted the distribution of velocity vectors for adhesin movement across the cell surface both for the adhesin shown in Fig. 1b,c,e (Fig. 1f, upper left) and for adhesins in seven other cells (one of which is shown in Fig. 1f, lower left). All these adhesins exhibited velocities predominantly in the upper-right and lower-left velocity quadrants of the plot, corresponding to the positive gradients of the adhesin trajectories in the Unrolled plot (Fig 1e) and to left-handed (LH) helical paths, as previously reported^16,19^. However, our higher resolution data reveal that adhesin movements deviate considerably from uniform LH helical paths and constant speed (Fig. 1f, left hand panels), which would have appeared as straight lines in Unrolled plots and as single points in the velocity distributions (see Fig. S1). Some cells showed an additional peak at the origin in the velocity distributions, corresponding to molecules that were stopped for part of the trajectory (an example is shown in Fig. 1f, lower left). For each of the 8 cells analysed, the peaks in the velocity distribution were taken and a linear fit applied to data from all 8 cells to establish the relationship between the velocity around the circumference and the velocity along the long axis (Fig. 1f, solid grey line). The resulting fit indicates that the adhesins move at an average angle α = 13.7 ± 1.8° to the cell long axis across a range of adhesin speeds up to ∼1.5 m s^-1^. This angle is consistent with the helical angle of 19.0 ± 11.5° previously reported by Nakane and co-workers for adhesin movements in gliding cells when tracked in 2D using antibody labelling^16^. A further study using 3D tracking of gold nano-spheres attached to SprB has reported that SprB moves with a helical pitch (distance along the cell long axis for one helical turn) P = 5.9 ± 0.7 μm^17^. Given a typical cell radius of ∼0.25 μm this corresponds to α between 13.4 and 16.8° consistent with our data and that of Nakane and co-workers^16^. However, it was reported that the gold nano-spheres move in right-handed (RH) helical trajectories contrary to the LH helical trajectories determined in our observations and those of Nakane and co-workers^16,19^. Such discrepancies in handedness can arise from errors in accounting for all reflections introduced in imaging, display, and image-processing. We took great care to avoid such errors (Methods, Supplementary Text 1) and confirmed that our methodology determined the correct handedness when test samples were moved in a known direction. Thus, our analysis confirms that SprB adhesins move in LH helices across the surface of *F. johnsoniae* in agreement with the observations of Nakane and co-workers^16,19^.

### Observed adhesin trajectories cannot be explained by a single closed loop model

The single-circuit closed loop model for the organisation of the gliding tracks predicts that an adhesin will follow the same path on each circuit of the cell (Fig. 1a). We identified trajectories from our datasets where the adhesin could be followed for more than one circuit before photobleaching. On rare occasions, the adhesin retraced a path very close to the one followed on the initial circuit of the cell, as predicted by the single closed loop model. Fig. 1b-e shows one example of this, where the terminated second circuit stays close to the path of the first circuit, albeit only barely within the estimated fluorophore localization precision. An example of more typical behaviour is shown in Fig. 2 (Video S3) where a single SprB molecule is tracked through nearly 4 circuits of the cell. In this cell the paths followed on subsequent circuits sometimes pass very close to those of previous circuits, but more often they do not. Notably, at some points the paths of the adhesin on successive circuits actually cross each other (e.g. L4 across L3 at X = 1.2 and 2.3 µm, L2 across L1, L3 and L4 between X = 3 and 4 µm). We also see paths that converge (e.g. R1 and R2 at X = 1.2). We considered the possibility that the changes in path between successive circuits in this and similar cells could arise from rolling of the cell (or of the gliding machinery) around the cell long axis during imaging. Numbering the paths in this particular cell in time order we see that rightward paths (well separated at X = 0.9 μm) are displaced from the top of the Unrolled plot in the order R4-R2-R1-R3 whilst leftward paths have the displacement order L3-L4-L1-L2 (assessed at X = 3 μm). Neither displacement sequence follows time-order, as would have been expected if unidirectional rolling had taken place. Furthermore, the sequences should be the same for rightwards and leftwards tracks if they were experiencing the same rolling motion (of any type), but again this is not what is observed. To provide additional confirmation that cells do not move under our experimental conditions we demonstrated that a protein that is known to be immobile in the outer membrane (SprE-HaloTag^34^) remains stationary when imaged (Fig. S5). Summarizing these experiments, our data indicate that adhesins do not normally repeat their trajectories on successive circuits of the cell, in contrast to the expectation of the single circuit closed loop model.

**Figure 2.**
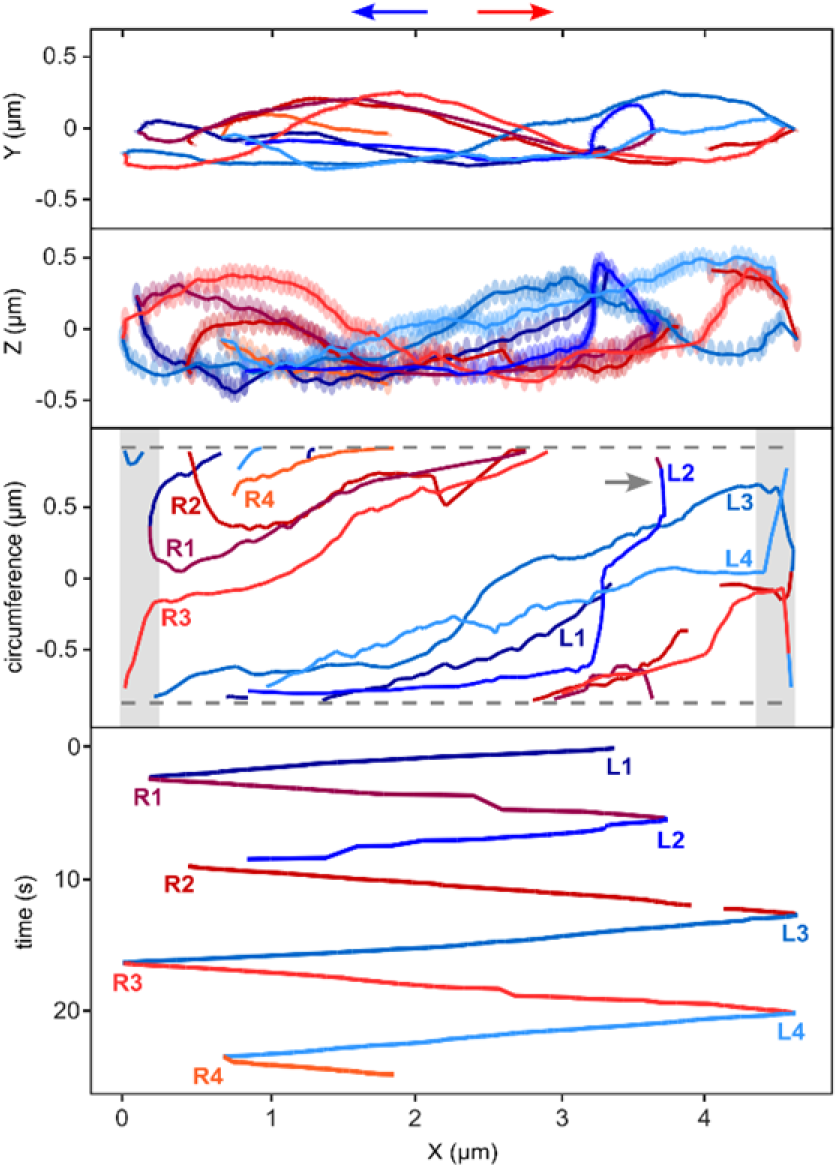
Adhesin trajectories cannot be explained by the single closed loop model. An SprB adhesin trajectory displayed as in Fig. 1e that takes different paths on successive circuits of the cell. Each pass of the cell length is coloured and labelled either L (leftward, shades of blue) or R (rightward, shades of red) and numbered in time order. An example of a mid-cell reversal, between R1 and L2, is indicated by a grey arrow in the Unrolled cell plot. See also Video S3.

An additional prediction of the single circuit closed loop model is that there can only be one helical path in each direction at any point along the cell length (Fig. 1a). However, we found that cells commonly had multiple adhesins moving in the same direction at the same time on separate paths. For example, in the cell shown in Fig. 3a (Video S4) pairs of adhesins (1&2, 3&4) simultaneously moved in the same direction on non-identical, non-overlapping trajectories across the same section of the cell body. These observations of multiple adhesin trajectories in parallel are incompatible with a single-circuit closed loop.

**Figure 3.**
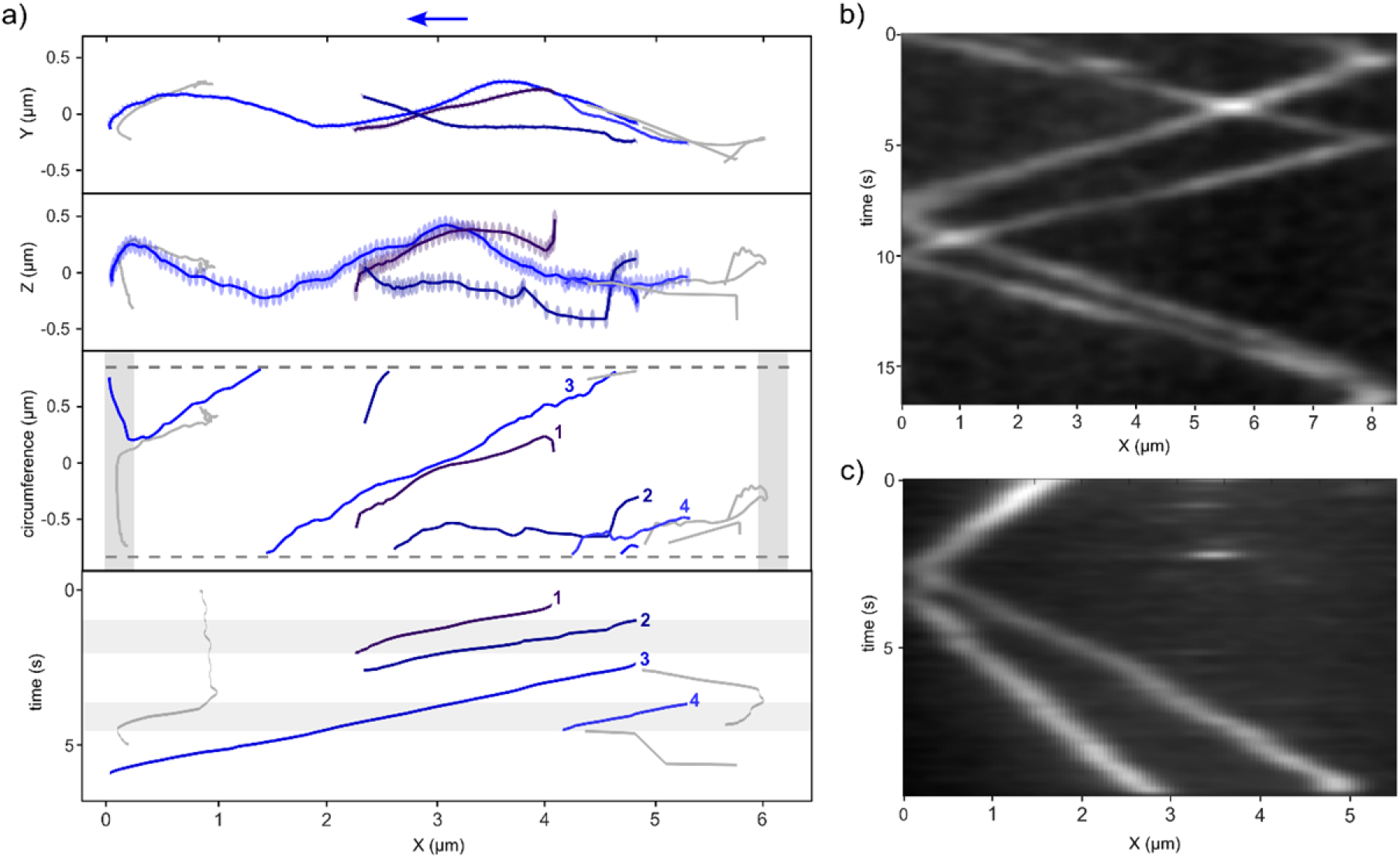
Adhesins in the same cell can simultaneously show different behaviours. (a) Adhesins can simultaneously move in the same direction on separate trajectories across the same section of the cell body. SprB adhesin trajectories displayed as in Fig. 1e. Selected leftward trajectories are highlighted in shades of blue and labelled 1 to 4 in order of appearance. Grey shading in the kymograph indicates the time periods within which two adhesins were simultaneously visible and moving in the same direction along non-overlapping paths. Note that trajectories starting after t = 0 arise when earlier localizations of the same molecule failed, most likely due to overlapping with other molecules. See also Video S4. (b,c) Adhesins in the same cell can move at different speeds. Kymographs obtained without single-molecule localization for 2 different cells. Each cell contains two distinct fluorescently-labelled adhesin molecules that travel simultaneously at different speeds. In **c** the two adhesins have overlapping point-spread functions for the first 2.5 seconds. See also Videos S5, 6.

A further prediction of the single circuit closed loop model is that all adhesin molecules will move at the same velocity at all times, because all are fixed to the same moving track (Fig. 1a). However, we observed multiple instances where two molecules moving simultaneously in the same cell travelled at distinctly different speeds (Fig. 3b,c). Often, the difference in speed between the two adhesins was maintained over multiple passes of the cell body. For example, in Fig. 3b (Video S5) the faster adhesin maintained an average speed of ∼1.6 μm s^-1^ and the slower adhesin ∼1.1 μm s^-1^ across 1.5 circuits of the cell. Conversely, in Fig. 3c (Video S6), two adhesins initially moved at approximately the same speed and then one adhesin accelerated whilst the other slowed. The observation that different adhesins can simultaneously move at different speeds indicates that multiple separate tracks are present, at least in some cells.

### Changes of adhesin direction far from the cell poles

In current gliding models, the adhesin trajectories are assumed to turn (change their long axis direction) only within the hemispherical end caps at each cell pole. Although this is normally the case, we also observed instances where adhesin trajectories turned several µm away from either pole. These mid-cell turns fall into two categories. Smooth turns have trajectories that resemble a turn in the end cap but occurring away from the pole (e.g. Fig. 2 R1 into L2, indicated by a grey arrow in Unrolled plot). By contrast, during sharp reversals (Fig. 4a) adhesin molecules abruptly change direction to approximately retrace their previous path briefly before diverging onto different paths. In some instances of sharp reversals, other adhesin molecules were observed simultaneously on the same cell, none of which showed any discontinuity in motion at the time of the reversal.

**Figure 4.**
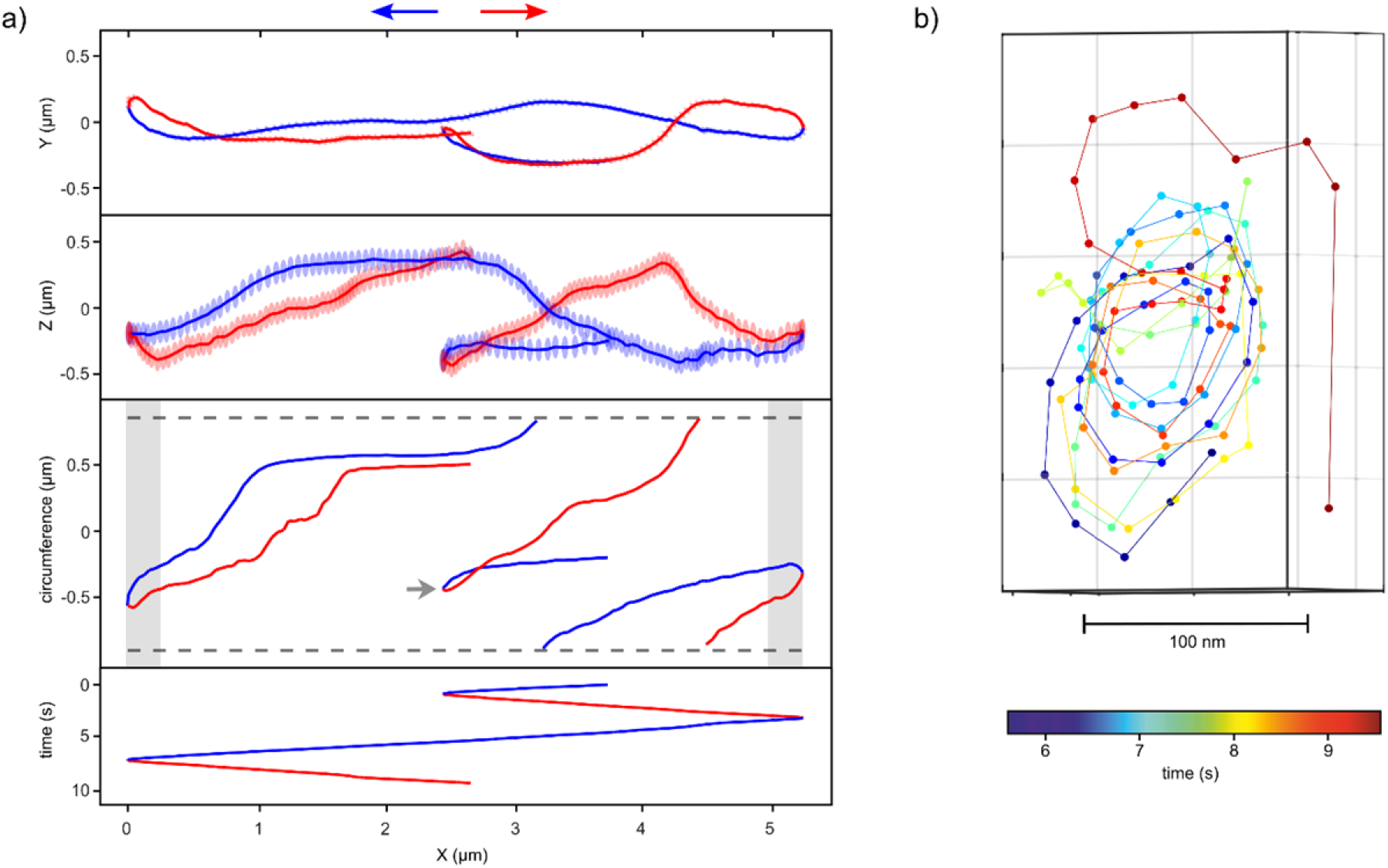
Trajectories of adhesins making sharp mid-cell turns or loops. (a) Mid cell turn by sharp reversal. An adhesin trajectory plotted as in Fig. 1e with the reversal highlighted by a grey arrow in the Unrolled plot. (b) A mini-loop trajectory. The projection is perpendicular to the local cell surface and viewed from outside the cell. In this representation the adhesin appears to move clockwise in a closed loop of diameter ∼100 nm. However, in reality the loop is ∼250 nm in diameter due to the 7-point smoothing window applied in the super-resolution tracking that reduces the diameter of a circular trajectory of this angular frequency (∼3 s^-1^) and frame rate (∼30 s^-1^) by a factor of 0.42. Only the last 11 of 17 rotations are shown, as other overlapping fluorescent molecules prevented tracking of the adhesin during the first 6 revolutions. Eventually the adhesin escapes the loop. Colours indicate time in accordance with corresponding Video S7.

### Mini-loops

In addition to mid-cell turns, we observed 2 trajectories in which SprB molecules made multiple rotations around a small circle without moving helically between the cell poles. The adhesins moved around both of these ‘mini-loops’ at ∼2 µm s^-1^, which is approximately the same speed as helically moving adhesins. Fig. 4b (Video S7) shows one of these mini-loops, which consisted of 17 revolutions at ∼3 revs/s around a loop ∼250 nm wide. Shrivastava and co-workers previously observed that cells subject to a shearing procedure were able to rotate adhesin-targeted polystyrene beads in ∼300 nm diameter circles ^21^ and it is possible that this phenomenon corresponds to the adhesin mini-loops described here.

### Adhesin trajectories cross and change direction in the cell end caps

The analysis described so far only characterizes how adhesins move along the cell body. We used 3D imaging to extend our analysis of adhesin trajectories into the cell end caps. 3D imaging is required for this task because adhesins moving in the end-caps have one main axis of motion (z) parallel to the optical axis. To maximize the number of trajectories observed across each end cap, we took advantage of cells where the field of illumination only covered a region within a few µm of a cell pole, thereby avoiding photobleaching the labelled adhesin molecules while they were in other parts of the cell. We describe localisations within the end cap using spherical polar co-ordinates (r′, θ, φ) with a centre given by the best-fit of all end cap data to a hemispherical surface (see Methods for more details).

Figs. 5a,b (Videos S8, 9) show azimuthal equidistant projections of all adhesin trajectories across two representative cell end caps, together with kymographs of the same data to indicate the timing of the trajectories. Adhesin molecules take a variety of paths across the end caps. Some trajectories make sharp turns in the end caps (e.g. trajectories 1, 2, 4 in Fig. 5b; Video S9). Some adhesins enter the end cap in near-identical locations but their trajectories subsequently diverge (e.g. trajectories 2, 3, 8 in Fig. 5a; Video S8). Trajectory 9 in Fig. 5a paused for ∼0.38 s and ∼0.72 s between periods of directional movement, indicating either that the adhesin detached from the underlying track or that the track itself stalled repeatedly at the pole. Similarly, trajectory 1 shows an adhesin that entered the pole moving directionally but then appears to transition to diffusive movement, suggesting detachment from the track complex. We observed a considerable range of adhesin speeds across the end cap, evident from the differences in spacing between data points for different trajectories. Many trajectories maintained consistently different speeds from other trajectories across the same end cap. For example, compare trajectory 5 in Fig. 5b (slow) and trajectory 2 in Fig. 5b (fast). Other trajectories appear to change speed after stopping (e.g. trajectory 9 in Fig. 5a) or slowing (e.g. trajectory 2 in Fig. 5a) near the pole.

**Figure 5.**
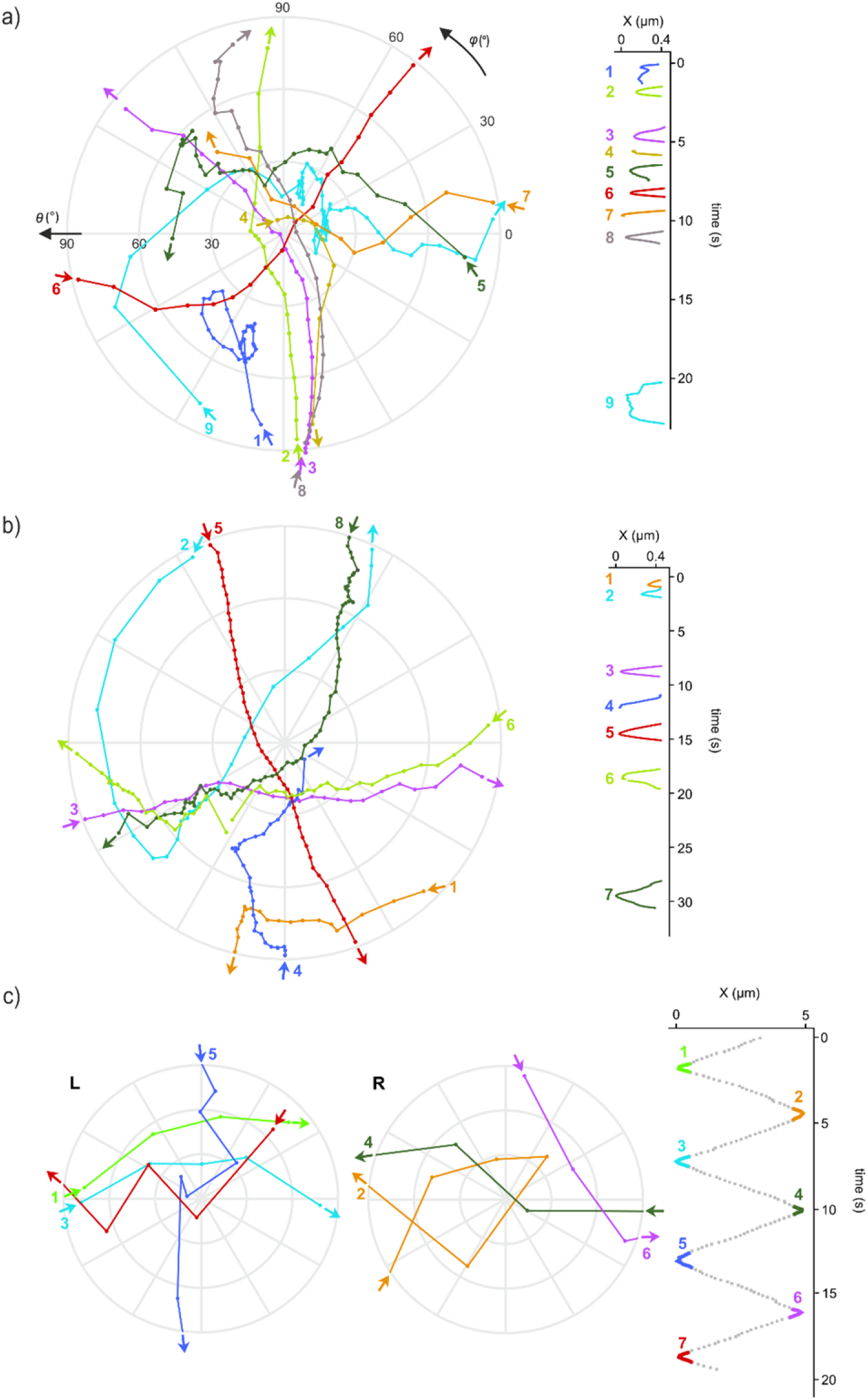
Adhesin trajectories across the cell end caps. (a,b) Multiple trajectories of adhesin molecules in one end cap of selected cells. (Left) Azimuthal equidistant projection of the end cap centred on the pole. Different trajectories are marked by numbers and different colours, and later trajectories overlay earlier ones. The direction of adhesin movement is indicated by start and end arrows for each trajectory. (Right) Adhesin distance X from the pole versus time, showing the time order of the trajectories. See also Videos S8, 9. (c) Fragments of the trajectory of a single adhesin molecule showing paths across the two end caps of the cell, designated left (L) and right (R). Each trajectory fragment is highlighted in the corresponding colour on the full kymograph on the right. This adhesin trajectory was acquired at a lower frame rate (10 fps) than the cells in (a,b). See also Extended Data 1 and Videos S10, 12.

Most strikingly, many of the trajectories cross one another in short succession. For example, in Fig. 5a trajectories 3 and 8 cross trajectories 5, 6, and 7 that occur in the intervening time. In this particular case the adhesin responsible for trajectory 8 is likely to be a different molecule from those travelling on the trajectories that it crosses, because trajectory 8 adhesin would not have had time to make a circuit of the cell and return to the crossing point in the time that has elapsed between the trajectories (5 s or less). To determine whether an adhesin might, nevertheless, cross its own earlier trajectory within the end cap, we sought to obtain very long, unbroken adhesin trajectories in cells with only one adhesin labelled. The duration for which we can observe the movement of a single adhesin is limited by photobleaching of the attached fluorophore. To extend the fluorophore lifetime we reduced the laser power with which the field of view was illuminated while increasing the exposure time to 100 ms in order to retain an acceptable signal-to-noise ratio, though at the expense of reduced temporal resolution. This allowed us to unambiguously track adhesins across multiple passes of the cell body after all others had photobleached, the most long-lived example of which is shown in Extended Data 1 (Video S12). This adhesin crosses its own trajectory multiple times in the end caps (Fig. 5c).

### A new open filament model for the organization of the gliding tracks

In this work we have used tracking of individual adhesin molecules to characterize the *F. johnsoniae* gliding track architecture at an unprecedented level of detail. This approach has revealed multiple new features of the adhesin trajectories that are inconsistent with current models for the mechanism of gliding.

First, the paths of individual adhesins can deviate substantially from ideal helical trajectories (Fig. 1f). Second, adhesin molecules do not normally retrace the same path on subsequent circuits of the cell but instead follow either partially or entirely different paths (Fig. 2). Third, when multiple adhesin molecules are tracked in the same cell, they most often travel on different paths rather than following the same path (Fig. 3a). Fourth, adhesin trajectories frequently cross each other, both in the cell end caps (Fig. 5) and along the cell body (Fig. 2, Fig. 3a). Fifth, simultaneously moving adhesins can move at different speeds (Fig. 3b,c). This phenomenon has also been reported for adhesins labelled with beads^23^ or antibodies^19^. However, these studies could not exclude the possibility that the differences in adhesin speeds were influenced by the multivalent and/or bulky nature of the labels. Sixth, we see instances where adhesins appear to reverse their direction (Fig. 2, Fig. 4a). Apparent adhesin reversals have also been reported using bead and antibody labelling^19,23^. However, those studies lacked the resolution to determine whether the adhesins were reversing back along a similar path or were on loops that turned before the end of the cell, two different reversing behaviours that are distinguished in the current work. In addition, these earlier studies could not exclude transfer of the polyvalent label between separate adhesins on separate tracks.

These behaviours of moving adhesins are incompatible with the current track model in which a single helical loop closes upon itself after exactly one circuit of the cell. If all adhesins were attached to a single closed loop then their trajectories would always repeat exactly, different adhesins would not follow distinct paths or move at different speeds, and their trajectories would not cross or reverse. Various modifications of the single closed loop model can individually accommodate several of the newly-described features of the gliding track: that the loop closes only after multiple circuits of the cell; that multiple loops are present; that loops form a network containing junctions at which adhesins do not always take the same branch. Nevertheless, none of these is able to account for all of the new observations. Moreover, they all require the gliding tracks to physically cross at points where the adhesin trajectories intersect, which appears incompatible with the known molecular features of the tracks^25^. The tracks are attached to the outer membrane through the lipid anchors of their lipoprotein subunits GldJ and GldK^25,35^. If two tracks were to cross, one would need to be displaced away from the membrane to allow the other to pass underneath it. This would require the lipid anchors of the displaced (but still moving) track to be transiently pulled from the inner leaflet of the outer membrane into the aqueous periplasm, a scenario that is mechanistically implausible. Second, the adhesins are attached to the gliding track through a disulfide linkage between the track and the OM protein SprF to which the adhesin is bound^25^. Adhesins would be unable to detach from one track to allow it to pass underneath another track without this disulfide bond being broken. Yet, a variant of SprF that appears incapable of reversing the disulfide linkage to the track supports normal adhesin movement over multiple passes of the cell body^25^, indicating that adhesins do not become stuck at crossing points if the disulfide bond cannot be cleaved. In summary, the molecular evidence argues against the idea that intersections of adhesin trajectories correspond to physical crossings of the gliding tracks.

To explain how adhesin trajectories can intersect without requiring the underlying tracks to cross, we propose a new model for the organization of the gliding tracks in which the tracks are not closed loops but are instead discontinuous open filaments (Fig. 6a). In this open filament model, we propose that cells contain multiple distinct track filaments. These would most likely be of variable lengths, with some considerably shorter than the length of the cell (Fig. 6a). This arrangement would allow two adhesins to follow intersecting paths without the tracks to which they are connected physically crossing, provided that the tail end of one track filament passes the overlap point before the head of the other filament arrives. This model would explain why adhesins in the end caps often cross the recent trajectories of other adhesins without interruption (Fig. 5), while on other occasions collisions between filaments causes the sharp changes of trajectory or temporary stalling observed for other end cap trajectories, and mid-cell reversals such as that of Fig. 4a.

**Figure 6.**
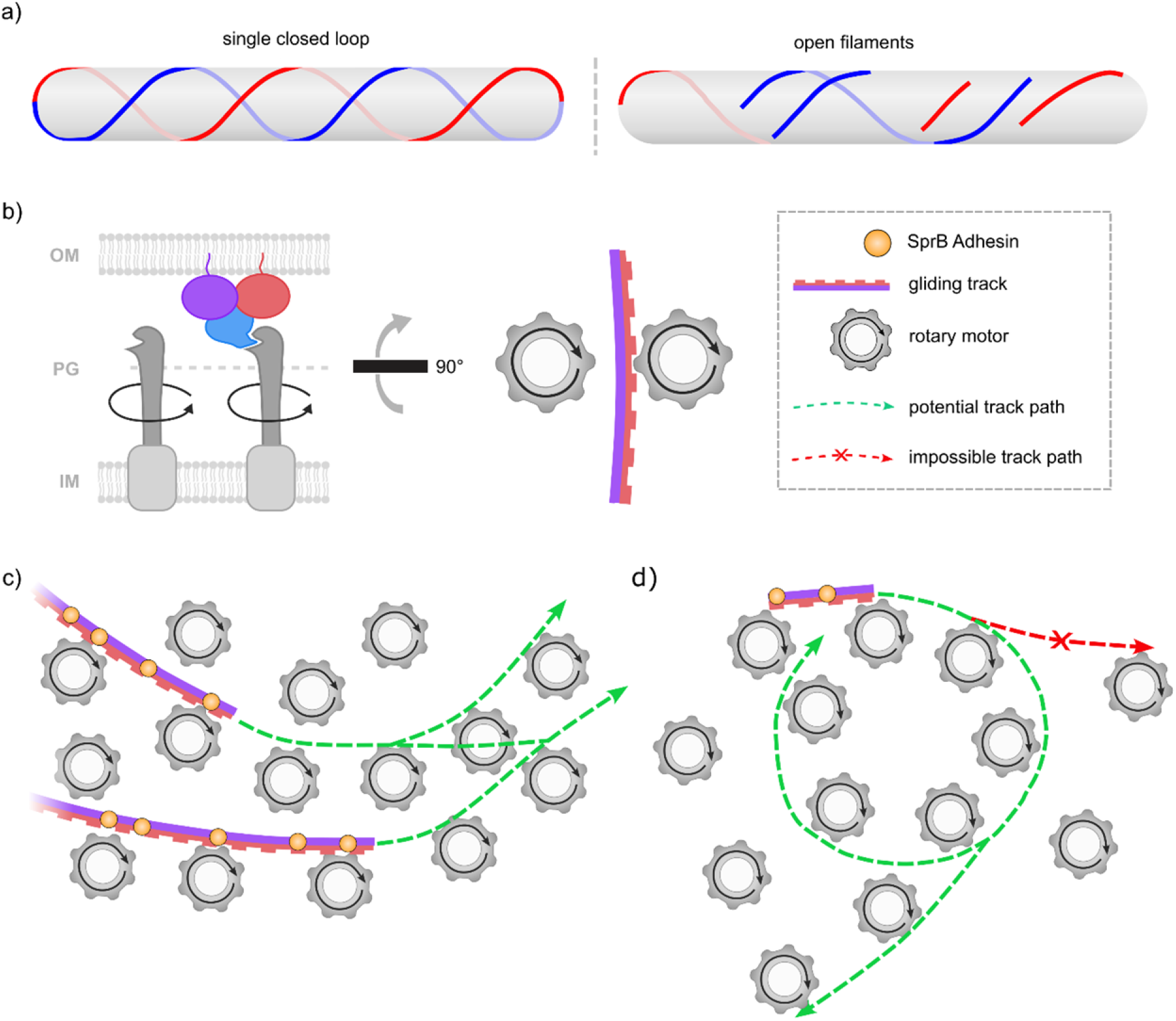
The open filament model. Schematic illustrations of track and adhesin behaviours within the open filament mode (a) Comparison of the current single closed loop model and the new open filament model. Track length, density and helical pitch are chosen for illustrative purposes. (b) The interaction between motors and track filaments. (Left) Schematic of the molecular organization. The rotating shafts of the motors can only interact with tracks in one filament orientation and tracks pass over motors with which they cannot interact. OM = outer membrane, PG = peptidoglycan cell wall, IM = inner membrane.(Right) Mechanical analogy viewed perpendicular to the membrane in which motors are depicted as cogs that interact with one side of the tracks. (c) The paths followed by distinct track filaments can converge or diverge as their leading ends interact probabilistically with different motor complexes. (d) The shortest tracks may fail to bridge between two motors on a canonical left-handed helical path, instead leading to a mid-cell reversal or mini-loop. (b-d) Illustrated components are not to scale, and directions of rotation are arbitrary.

Current structural evidence suggests that the gliding track is propelled by motor proteins that unidirectionally rotate in place^27,28^. Due to the asymmetric nature of the gliding track filament^25^ motors can only productively contact the track in one orientation (Fig. 6b). In our open filament model, the pathway by which the gliding filaments move between motor complexes is not fixed but instead depends on the probability of different motor complexes binding the leading end of an approaching filament (Fig. 6c). This stochasticity in filament pathways would explain our observations that adhesin trajectories can vary between successive circuits along the cell or converge and diverge from each other (Fig. 2), and deviate from a fixed helical pitch (Fig. 1f). When the shortest filaments are too short to span a gap between two motors on a canonical helical path (Fig. 6c) the open filament model predicts that mid-cell reversals (Fig. 2), and in extreme cases mini-loops (Fig 4b), can occur.

Substantial changes in adhesin pathway only occur a few times per cell circuit at most. The frequency of such events will depend on many factors including filament length and rigidity, the number of motors in contact with the filament at any one time, and the density and homogeneity of motor complex distribution. Adhesins attached to short filaments may be more likely to perform aberrant behaviours because these are in contact with fewer motors at any given time and therefore have more flexibility to change their path. This may be occurring for example in Fig. 2 where there are reversals not at the end caps. It is also possible that filaments will preferentially follow recently-used paths if the adhesin-carrier protein complexes find it easier to move through areas of the OM where the highly cohesive lipopolysaccharide molecules^36^ have already been pushed apart by previous adhesin passage. Filament movements may also be constrained if multiple filaments are aligning together in bundles as suggested by cryo-electron tomography^19^.

The open filament model naturally explains the observation that adhesins in the same cell can move simultaneously at different speeds, because separate filaments are not constrained to move at identical speeds. Speed differences might arise from factors including variations in the number of motors engaged by a filament and in resistance to filament movement, each of which might depend on filament length. Cases where two adhesins maintain different speeds over multiple passes of the same cell (Fig. 3b) indicate that speed of adhesin movement along the cell body is determined by the properties of individual filaments rather than the properties of the cell or the motors in any particular location.

The open filament model is also able to address previously un-appreciated problems faced by closed loop models that occur as the cell grows. A closed loop of constant helical pitch would need to lengthen as the cell elongates. However, because it is a sealed loop it has no obvious way of inserting additional track subunits. By contrast, there is no necessity for open filaments to change length as the cell expands. Filament elongation, if required, is straightforward for an open filament because new subunits can be added at either of its two free ends. During cell division, a closed loop track in the mother cell would need to be cut in two and the resulting fragments individually resealed to form new closed loops in the daughter cells. However, no such process is required if the tracks are already discontinuous. Both the gliding track and the gliding adhesins have been observed to accumulate at the division septum during cell division^5,35^. This is as expected under an open filament model, because track filaments moving along the outer membrane are unable to pass the division septum where the outer and inner membranes are no longer directly juxtaposed. By contrast, such accumulation would not be predicted if a closed loop was simply being sliced in half. The closed loop model would therefore need to either invoke an additional track condensation process or acknowledge that the track can still move once fragmented, raising the question as to why it does not do so all the time.

In summary, the features of the Bacteroidota gliding track that are newly described here can be explained by a radically altered model in which the gliding track is composed of multiple open filaments rather than forming a closed loop as previously envisaged. This arrangement is strikingly similar to that of bacterial cytoskeletal systems based on the actin homologue MreB and the tubulin homologue FtsZ. Like the gliding filaments, these polymers were originally thought to form continuous loops but were later revealed by advances in imaging methodology to form mobile open filaments. MreB forms linear polymers that are on average less than 200 nm in length and are propelled around the cell circumference by the machinery that synthesises the cell wall^37-39^. FtsZ forms linear polymers involved in organising cell division. These filaments move around the division septum by a GTP-powered treadmilling mechanism in which protomers are added to one end of the filament and released from the other end^40,41^. The common observation of discontinuous filaments across these different systems suggests that this architecture may represent a general organising principle for mobile bacterial polymeric filaments.

## Methods

### Cells and Strains

*F. johnsoniae* strain AK_73 (UW101 Δ*porV halo-sprB*)^28^ was used in this study to allow fluorophore labelling of the SprB adhesin. AK_73 expresses SprB with a HaloTag domain inserted between the signal peptide and the N-terminus of the mature protein. In order to minimize proteolytic cleavage of the fusion protein by endogenous secreted proteases we employed a Δ*porV* genetic background which prevents export of most proteases through the Type 9 Secretion System.

To assess cell immobility in agarose and to determine the localization accuracy of single fluorescent molecules in our microscope we used strain FL_147 (UW101 *sprE-halo*)^34^ which expresses a SprE-HaloTag fusion protein that is stationary in the outer membrane^34^. To assess the specificity of HaloTag labelling we used strain FL_030 (UW101 Δ*porV*)^28^ which has no HaloTag on SprB.

### Cell preparation for microscopy

Cells were grown overnight in Casitone Yeast Extract (CYE) medium^42^ at 28 °C with 185 rpm shaking. The cells were then diluted to an OD600 of 0.05 in Motility Medium^43^ and cultured for a further 3 hours. 1 µl of cells was incubated with 1 nM Janelia Fluor 646 HaloTag ligand (Promega) for 30 minutes. Cells were harvested by centrifugation at 8,000 *g* for three minutes, washed three times in PY2 medium^44^ and resuspended in PY2 medium. For 3D imaging cells were spotted onto 1% agarose PY2 agarose pads formed in 65 µl Gene Frames (Thermo Fisher Scientific) and imaged through a coverslip.

### Parallax microscopy

Samples were imaged in a custom-built inverted microscope comprising a Nikon 1.45 NA Plan Apo TIRF Oil immersion objective with Cargille Type B immersion oil, dichroic mirror Dio02 r635 (Semrock) and emission filter LP635 (Semrock). The microscope was fitted with both a course stage and a piezoelectric stage (P-517.3CD, range 100 µm in *x* and *y*, 20 µm in *z*, PI (Physik Instrumente) Ltd). A field of view 8.5 µm wide (full width at half maximum) was illuminated at an average power of 5,200 W cm^-2^ with a 633 nm Helium-Neon laser (Melles-Griot 25-LHP-991-230). We used a “D-mirror” (Thorlabs, BBD05-E02) in a conjugate back-focal-plane (bfp) to split the bfp into 2 equal halves. We used each half to form an image (a “view”) of the specimen plane, side-by-side, on an Electron-Multiplying CCD (EMCCD) camera (iXon Ultra 888, Andor), water cooled to -80°C. Except for the cell of Fig. 5c, images were acquired at full EM gain (300) with 30 ms exposures (34.494 ms cycle time) and a vertical shift speed of 4.33 µs. The final magnification was ∼200x, resulting in pixels measuring 51.5 nm square.

To calibrate axial localizations (*z*), we immobilised Crimson 200 nm FluoSpheres™ Carboxylate-Modified Microspheres (FF8806, ThermoFisher Scientific) on a coverslip and stepped through 1.008 µm in *z* (20 steps of 50.4 nm) with the EMCCD camera acquiring six images at each *z*. Fluorescent beads were localized (*x*,*y*) in each image (see ‘3D tracking’) and localizations averaged for each *z*. The separation between averaged localizations in the two views was fitted as a third order polynomial function of *z* to yield a calibration curve for *z*.

### 3D tracking

We identified single fluorescent molecules in each view in each frame of each saved video using ADEMS code^45,46^ as follows. For each candidate molecule, ADEMS calculates the background corrected signal-to-noise ratio (SNR) as the sum of pixel intensities within a circular region of radius 8 pixels centred on the candidate molecule, divided by the standard deviation of pixel intensities within a concentric 10 x 10 pixel region excluding the circular region. Within the ADEMS code, we excluded candidate molecules with SNR below 0.25-0.3 and molecules separated by < 2-3 pixels (that cannot be reliably tracked separately) to produce a candidate localisation list. We selected limits for each imaging session depending upon the density of candidate molecules per cell, with higher SNR and lower separation limits for more densely labelled cells.

Using custom-written code we defined trajectories in each view by linking a molecule localized in the current frame to the closest molecule within 300 nm in the previous frame (or failing that within 600 nm, 2 frames previously), only if its intensity was between 0.5-100x the previous intensity (occasionally 0.5-300x for cells with very dense JF646 labelling). We paired trajectories between the two views if they overlapped closely in time and the distance between them in their first shared frame was ≤ 15 pixels for low (or 7 pixels for high) density labelling. Where more than 2 trajectories could be paired as above, no pairing was assigned and a warning was sent to the user. When an ADEMS candidate in only one frame in only one of a pair of trajectories failed the SNR criteria (e.g. due to photo-blinking or shot noise; we used the minimum possible laser intensity to minimise photobleaching), we ‘restored’ it if it was within 15 (7) pixels of the paired localization. Otherwise we linked the paired trajectories across the gap, but did not attempt to fill the gap in. ADEMS typically returns a single localization at the centroid of two molecules which are very close in a single view. This created gaps several frames long when molecules coincide in 2D. We assigned localizations either side of such gaps to different trajectories.

Finally, we calculated 3D trajectories from paired view trajectories using the calibration curve for *z*, obtained using microspheres as described above. We estimated fluorophore localization uncertainties of [13, 34, 90] nm in the microscope co-ordinate frame [*x*, *y*, *z*], where *z* is the optical axis and *y* the direction of parallax splitting, using trajectories obtained as above for the immobile outer membrane protein SprE^34^ (Figure S5). For display we smoothed all trajectories with a 7-point running boxcar average filter. Co-ordinate transformations to the new (X, Y, Z) axes, where X is the long axis of the cell, rotate the uncertainty ellipsoid in (X, Y). We did not attempt to quantify the further uncertainty in the location of the point of contact between SprB and the gliding track due to the ∼150 nm contour length of SprB^16^ between that point and the SprB-distal location of the fluorophore. For this reason error-bars may underestimate the true localization uncertainties. Image analysis was performed using custom written MATLAB (Mathworks) scripts.

### Data selection and categorization

We acquired a total of 1100 videos containing ∼1600 immobilized *F. johnsoniae* cells with fluorescently labelled SprB. We manually selected and saved 572 videos containing ∼800 cells which showed fluorescent molecules that lasted more than several seconds before photobleaching. We manually selected 171 of these cells for further analysis according to the following criteria. 1. Images well focussed. 2. Images fully within the calibrated *z* range. 3. No detectable stage drift. 4. Contained single molecule trajectories ≥ 20 frames (0.69 s) in order to exclude possible non-specifically attached dye molecules (see Fig. S6). 5. Contained sufficient trajectories for convergence of a cylindrical fit to the cell body or a hemispherical fit to the end cap (see below), or showed mid-cell reversals, mini-loops or any other clear non-helical motions. We assigned 3D trajectories to individual cells identified by segmentation of parallax brightfield images.

For Figs 3b and 5c only, we obtained videos of ∼400 cells at 10 fps and a laser power of 2,200 W cm^-2^. The cell of Fig 5c (see also Extended Data 1 and Video S12) included the long-lived molecule shown in that figure.

### Unrolling cells - cylindrical co-ordinates

We used the set of all 3D localisations for each cell to fit the cell surface as a sequence of 11 linked cylindrical sections (i.e. each cell surface fitted as a tube that need not be straight). Cylindrical fits were obtained as follows (see Fig. S7). 1. We rotated images about the optical axis, (*x, y, z*) → (X′, Y′, Z = *z*), to align the cell long axis to the new X′-axis, by minimizing the sum over all localizations of squared distances from the (X′, *z*) plane. 2. We made an initial approximation to the cell long axis as a 3^rd^ order polynomial *Y′(X′)*, obtained by trust-region reflective least-squares fitting of 3D localizations projected onto the (*X′, Y′*) focal plane. 3. We divided the cell into 10 preliminary sections of equal length along the *X′*-axis and calculated the centre of each preliminary section as ***<u>S</u>i*** *= (X′, Y′, z)I* = (*X′i, Y′(X′i),* 0), with *X′I = X′min* + (*i* - ½)( *X′max - X′min*)/10 (*I* = 1 to 10) being the mid-points of each preliminary section, *X′min* and *X′max* the *X′* co-ordinates of left- and right-most localizations respectively. 4. We divided the cell into 11 sections using ***<u>S</u>i*** as ‘section points’ with the 2 terminal sections each half the length of the 9 other mid-cell sections. 5. We defined the boundaries of each mid-cell section as the planes normal to the *X′*-axis and containing the section points, assigned localizations between these boundaries to the corresponding section, and assigned remaining localizations to the nearer terminal section. 6. We transformed all localizations by a rotation in the (*X′, Y′*) plane that aligned a new local *X*-axis (*Xi*) to a line joining the 2 section points bounding the section containing the original localization, keeping the new local *Y*-axis (*Yi*) in the original (*X′, Y′*) plane, and retaining the original *z*-axis perpendicular to both *X* and *Y*. 7. We projected the resulting localizations into the new (*Yi, z*) plane and found the centre (*Y0, z0*) and radius *R* (in the range 180-350 nm) of the circle that minimized the cost function *Σj (rj-R)^2^*, where *j* indicates localizations in the section and *rj^2^ = (Yj-Y0)^2^ + (zj-z0)^2^*. 8. We transformed to coordinates with an origin at the centre of the circle fitted in step 7 and the cell midline along the new *X* axis, then converted the transformed localizations ***<u>R</u>ij*** *= (X, Y, z)ij* into local cylindrical co-ordinates *(X, r, φ)ij*. *X* was defined as the contour length along the polynomial fit *X′(Y′)*. “Unrolled” plots of Figs 1-3 and Extended Data 1 show localizations as (*X, Rφ*) such that *Rφ* in mid-cell sections represents distances around the circumference of the cell. In end cap sections, unrolling shows a distorted (Mercator) projection. These data are displayed under greyed shading in Figs 1-3.

### Spherical co-ordinates for cell poles

Within each cell end cap we converted the localizations into local hemispherical co-ordinates (*r, θ, φ*), with *θ* running from 0 at the cell pole to 90° at the equator of the hemisphere, and *φ* matching the cylindrical *φ* co-ordinate (Figure S7). The hemisphere centre ***<u>S</u>*** was determined as follows. Steps 1 and 2 were performed as described above for cylindrical transformations. 3. An initial estimate for the cell pole was made from the polynomial fit Y′*(X′)*, and for ***<u>S</u>*** as a point on the cell long axis at a distance *R0* = 250 nm from the pole. 4. We refined the location of ***<u>S</u>*** by exhaustive minimization over a set of points spaced by 1 nm in all three axes of a cost function *f = Σj (rj-R0)^2^*, where *j* indicates localizations with the end cap (that is, points with X’ between ***<u>S</u>*** and the cell extremum) and *rj^2^ = (xj-Sx)^2^ + (yj-Sy)^2^ + (zj-Sz)^2^*. 5. We repeated step 4 with refined sets of end cap points, until the fit converged to 0.5 nm in all directions. For end caps where this procedure failed due to sparsity of data-points, we chose ***<u>S</u>*** by hand to minimize |*rj-R0*| and to place ***<u>S</u>*** close to the apparent cell axis as judged by points in the end cap and nearby. End cap plots (Fig. 5) show azimuthal equidistant projections of localizations in hemispherical co-ordinates, with no attempt to represent the localization uncertainties.

## Extended Data

**Extended Data 1.**
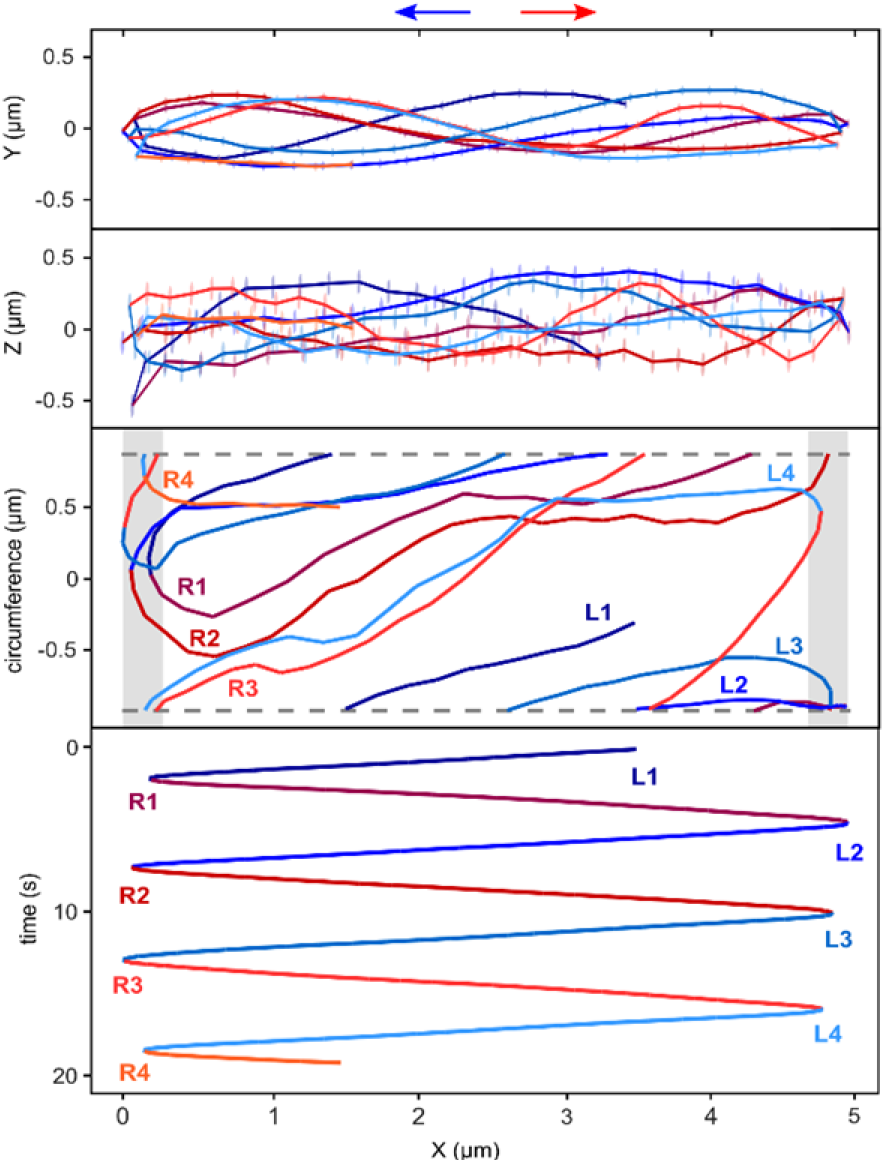
A single adhesin trajectory displayed as in Figure 1e, for the cell of Figure 5c. These data were acquired at a lower frame rate (10 fps) compared to the rest of the data in this publication. This adhesin confirmed the observations of Fig. 2: it followed distinct paths on each successive pass of the cell body with some paths in the same direction crossing over one another (e.g. R2 and R3, L2 and L3), and others totally non-overlapping (e.g. R1 and R2, L3 and L4).

## Statistics and reproducibility

Data for this paper were acquired across more than three biological repeats to ensure that the adhesin behaviour observed was representative. All adhesin behaviours presented in this paper were observed across multiple biological repeats. Controls for non-specific dye binding and behaviour of stationary SprE were acquired across three biological repeats.

## Data availability

All relevant data are included in the article and/or supporting information. Original code used in this study is available at https://zenodo.org/records/22712984.

## Acknowledgments

RMB, HLM and SC were supported by EPSRC Established Career Fellowship EP/S036660/1 (RMB). RLJ and BCB were supported by European Research Council Advanced Award 833713 (BCB) and Wellcome Trust Discovery Award 317933/Z/24/Z (BCB). RLJ was additionally supported by Medical Research Council studentship MR/N013468/1. AK was supported by a Biotechnology and Biological Sciences Research Council studentship. We thank Rut Carballido-Lopez for discussions and Ren Lim for the gift of LabVIEW code.

## Author Contributions

HLM built the Parallax microscope. HLM, RLJ, AK. and SC collected data. HLM, SC, and RLJ wrote code. HLM, RLJ, SC, and RMB analysed the data. Visualisation was by HLM, RLJ, SC, and RMB. The open filament model for the gliding track was conceived by RLJ. The manuscript was written by HLM, RLJ, BCB, and RMB, and all authors agreed on the final version. BCB and RMB conceived and supervised the project and secured funding.

