## Supplementary Information for "Open filament tracks drive Bacteroidota gliding motility"

**Video S1**

Immobilized cell of Figure 1b-e, illustrating how depth is encoded as differential movement in the parallax shifted (vertical) axis. One molecule travels up and down the length of the cell, with depth changes encoded in the vertical separation between red and cyan images. Note also the variation of the point-spread-function with depth. The other molecule is immobile, and photobleaches approximately halfway through the video. Images are smoothed with a Fourier Bandpass filter from 6-100 pixels and the grey-scale linearly stretched, for easy visualization of molecules.

**Video S2**

The same data as Video S1 of a fluorophore-labelled SprB adhesin molecule moving in the immobilized cell of Figure 1b-e, illustrating how 3D trajectories are obtained from the parallax images. Raw images.

**Video S3**

Overlaid parallax views as in Video S1, for the cell of Figure 2. Initially there are 2 fluorescent adhesins moving in the cell. After one of these photobleaches (~23% of the way through the video) the remaining adhesin, tracked in Figure 2a, makes a further ~3 full circuits of the cell. Images are smoothed with a Fourier Bandpass filter from 6-100 pixels and the grey-scale linearly stretched, for easy visualization of molecules.

**Video S4**

Overlaid parallax views as in Video S1, for the cell of Figure 3a. 2 cells are visible in this video, each labelled by many fluorescent adhesins and outlined by these and by autofluorescence in the early frames. The lower cell spanning the diagonal of the video is the source of the trajectories shown in Figure 3a. After progressive photobleaching single adhesins can be resolved and tracked. Images are smoothed with a Fourier Bandpass filter from 6-100 pixels and the grey-scale linearly stretched, for easy visualization of molecules.

**Video S5**

Overlaid parallax views as in Video S1, for the cell of Figure 3b. Two distinct adhesins start at different points along the cell body. The adhesin that starts in the bottom-left moves faster than the adhesin that starts in the top-right, and after multiple passes of the cell body the former adhesin catches up to the latter. These data were acquired at a lower frame rate (10 fps) compared to the rest of the data in this publication. Images are smoothed with a Fourier Bandpass filter from 5-70 pixels and the grey-scale linearly stretched, for easy visualization of molecules.

**Video S6**

Overlaid parallax views as in Video S1, for the cell of Figure 3c. For the first 2.5 seconds, 2 adhesins travel together from mid cell over the left-most pole at approximately the same speed (~0.6 μm $s^{-1}$), unresolved as a bright single fluorescent spot. After changing their long-axis direction at the cell pole, both adhesins traversed the same section of the cell within a few seconds of each other, in the same direction, but on different paths. The faster adhesin moved at ~0.8 μm $s^{-1}$, the slower adhesin appeared to reverse its direction and travel a short distance back to the cell pole before resuming its rightward traversal of the cell length at a reduced speed of ~0.5 μm $s^{-1}$. Images are smoothed with a Fourier Bandpass filter from 6-100 pixels and the grey-scale linearly stretched, for easy visualization of molecules.

**Video S7**

Overlaid parallax views as in Video S1, for the cell of Figure 4b. The molecule that was moving in a “mini-loop” is at the left pole of the lower cell, near the point where it touches the other cell. Images are smoothed with a Fourier Bandpass filter from 6-100 pixels and the grey-scale linearly stretched, for easy visualization of molecules.

**Video S8**

Animation of projections of the end-cap trajectories of Figure 5a. New localizations in each successive frame are added sequentially in colours corresponding to each trajectory, over grey dots showing all localizations in all frames. Azimuthal equidistant and each of 3 Cartesian projections are shown, as well as long axis position ($x$) versus time. Trajectories follow many different paths. Several trajectories cross the paths of earlier and later trajectories.

**Video S9**

Animation of projections of the end-cap trajectories of Figure 5b, as in Video S8. Two trajectories make sharp turns in the end-cap.

**Video S10**

Animation of projections of the left-hand end-cap trajectories of Figure 5c, as in Video S8. The data for Figure 5c were acquired at a lower frame rate (10 fps) compared to the rest of the data in this publication (see also Extended Data 1). The animation includes all trajectories obtained for this cell. In Figure 5c, only data later than 17 s are shown, after all but one adhesin label had photobleached. This adhesin likely made ~6.6 full circuits of the cell, and was the only labelled adhesin for the last ~3.5 circuits shown in Figure 5c. Thus, the last 7 end-cap transits, shown in Figure 5c, are all certainly made by a single adhesin.

**Video S11**

Animation of projections of the right-hand end-cap trajectories of Figure 5c, as in Video S10.

**Video S12**

Overlaid parallax views as in Video S1, for the cell of Figure 5c (see also Extended Data 1). These data were acquired at a lower frame rate (10 fps) compared to the rest of the data in this publication. Only a single adhesin is labelled in this cell, which completes multiple circuits of the cell body before photobleaching. Images are smoothed with a Fourier Bandpass filter from 5-70 pixels and the grey-scale linearly stretched, for easy visualization of molecules.

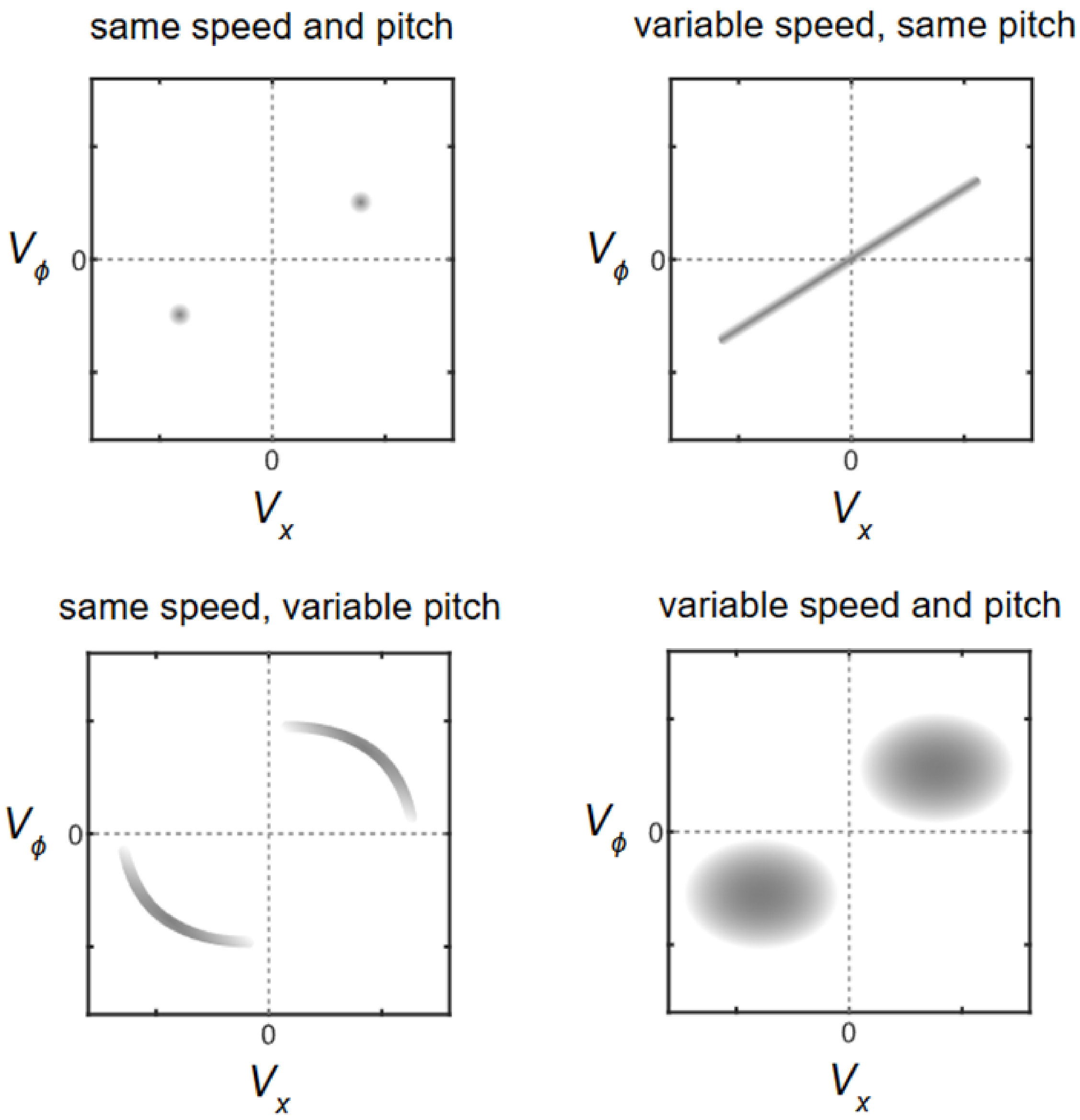


**Figure S1:** Schematic velocity plots to demonstrate the expected appearance given constant speed and/or helical pitch of a moving adhesin. All plots assume that the adhesins follow a left-handed helical trajectory.

**Supplementary Text 1**

We took great care to ensure that our 3D localization procedure did not introduce a parity reversal, which would cause left-handed helices to appear right-handed and *vice-versa*.

We established the correspondence between movements in the lab and movements on-screen in our two parallax viewpoint images. While recording images of a sample of 200 nm fluorescent beads attached to the cover glass, we rotated by hand each of the right-handed micrometer screws controlling our mechanical microscope stage, one at a time, clockwise as viewed from their pushing side. This moved the stage in the sequence: **away, left, up** as viewed by the user (Figure S2). Large stage movements in these directions were directly visible to the naked eye, confirming the directions unambiguously.

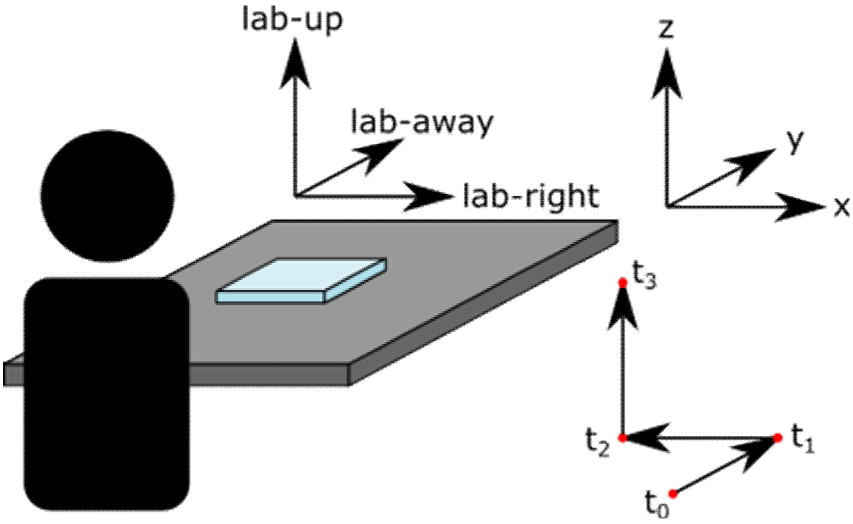


**Figure S2:** Schematic showing the sequence of movements of particles attached to the stage: away from the user ($t_0 \rightarrow t_1$), to the user's left ($t_1 \rightarrow t_2$) and finally up ($t_2 \rightarrow t_3$). The sequence was performed by manual adjustment, either of micrometer screws or the controls of a 3-axis piezo-electric stage. The co-ordinates of our recovered 3D localizations (x, y, z) are also indicated.

Figure S3 shows video frames at time-points $t_0$ to $t_3$ during such a sequence, performed by hand with the stage micrometer screws (to establish the directions, A) or a piezo-electric stage, to achieve smaller and slower movements suitable for our tracking algorithms, B). In both cases the movements on the screen are in the same directions, confirming that the small motions in the lab frame caused by the piezo stage were indeed in the directions shown.

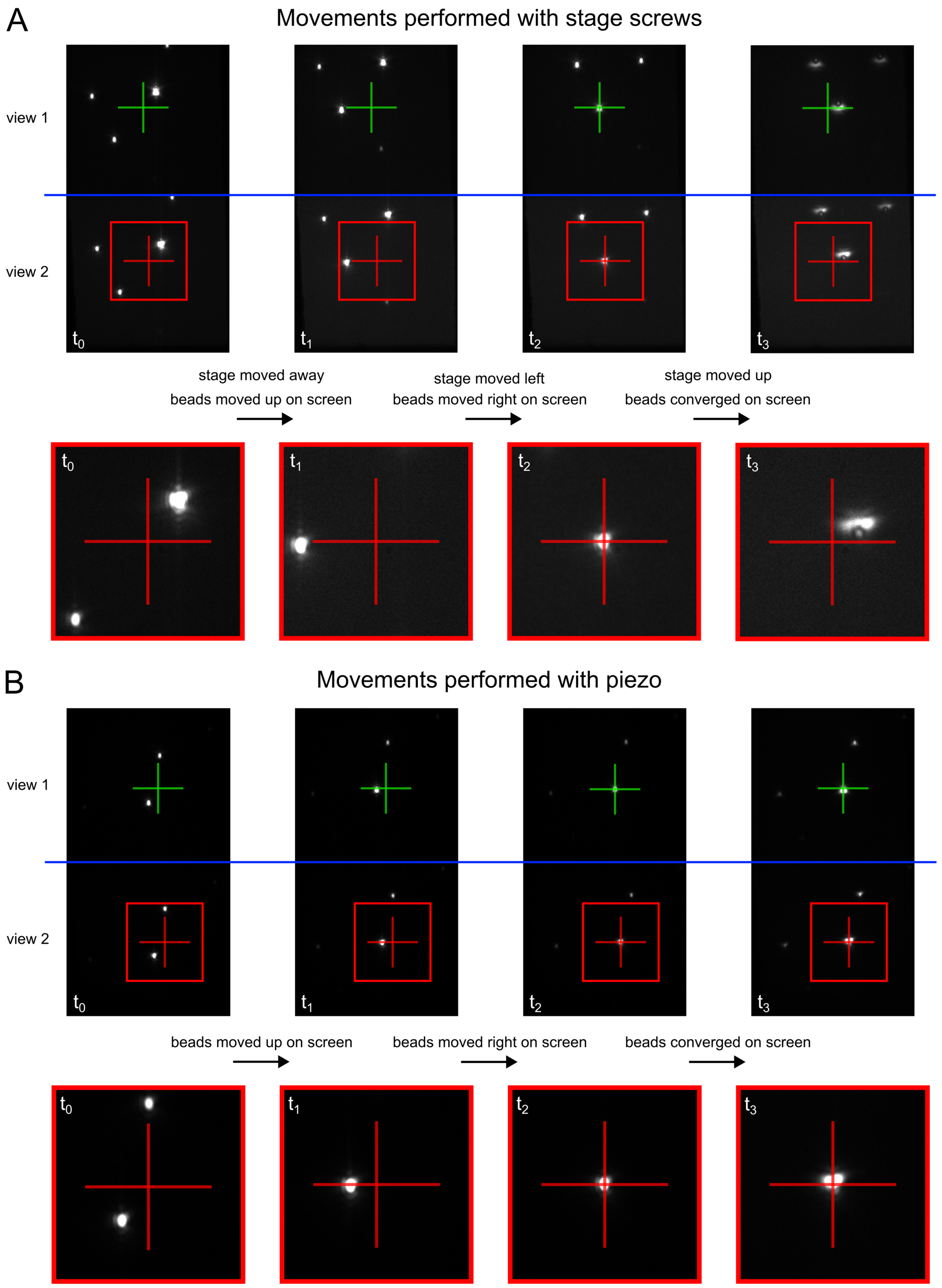


**Figure S3:** Video frames (upper) from recordings of the straight-line stage movements described in figure S2, with micrometer screws (A) or piezos (B). The directly observed lab-frame movements of the stage and the corresponding movements of bead images are indicated between the frames. Parts of each frame corresponding to the two parallax views are separated by a blue line. Cross-hairs fixed at the position of a particular bead in frame $t_2$, and enlarged views of the areas within the red squares (lower) are added to make it easier to see the bead movements.

Figure S4 displays 3D particle trajectories recovered from the videos illustrated in Figure S3B using the particle tracking algorithm described in the Methods section of the main text. From the viewpoint of the user, located in the **-y** direction and oriented as **+z** up, the movement is correctly rendered as **away, left, up.** This demonstrates that our recovered 3D locations, when displayed in a right-handed Cartesian axis system, preserve the parity of the real world. The correspondence between directions in the lab and the recovered 3D co-ordinates is shown in Figure S2.

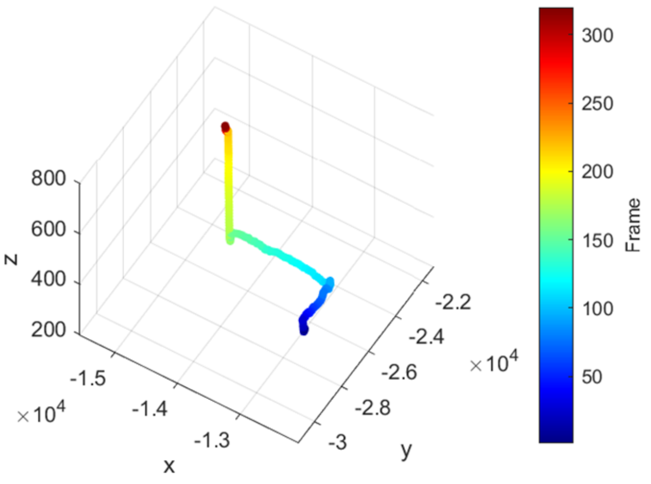


**Figure S4:** 3D particle trajectories recovered from the videos illustrated in Figure S3 using the particle tracking algorithm described in the methods section of the main text. The movement is correctly rendered as **away, left, up**, confirming that our recovered 3D locations preserve the parity of the real world.

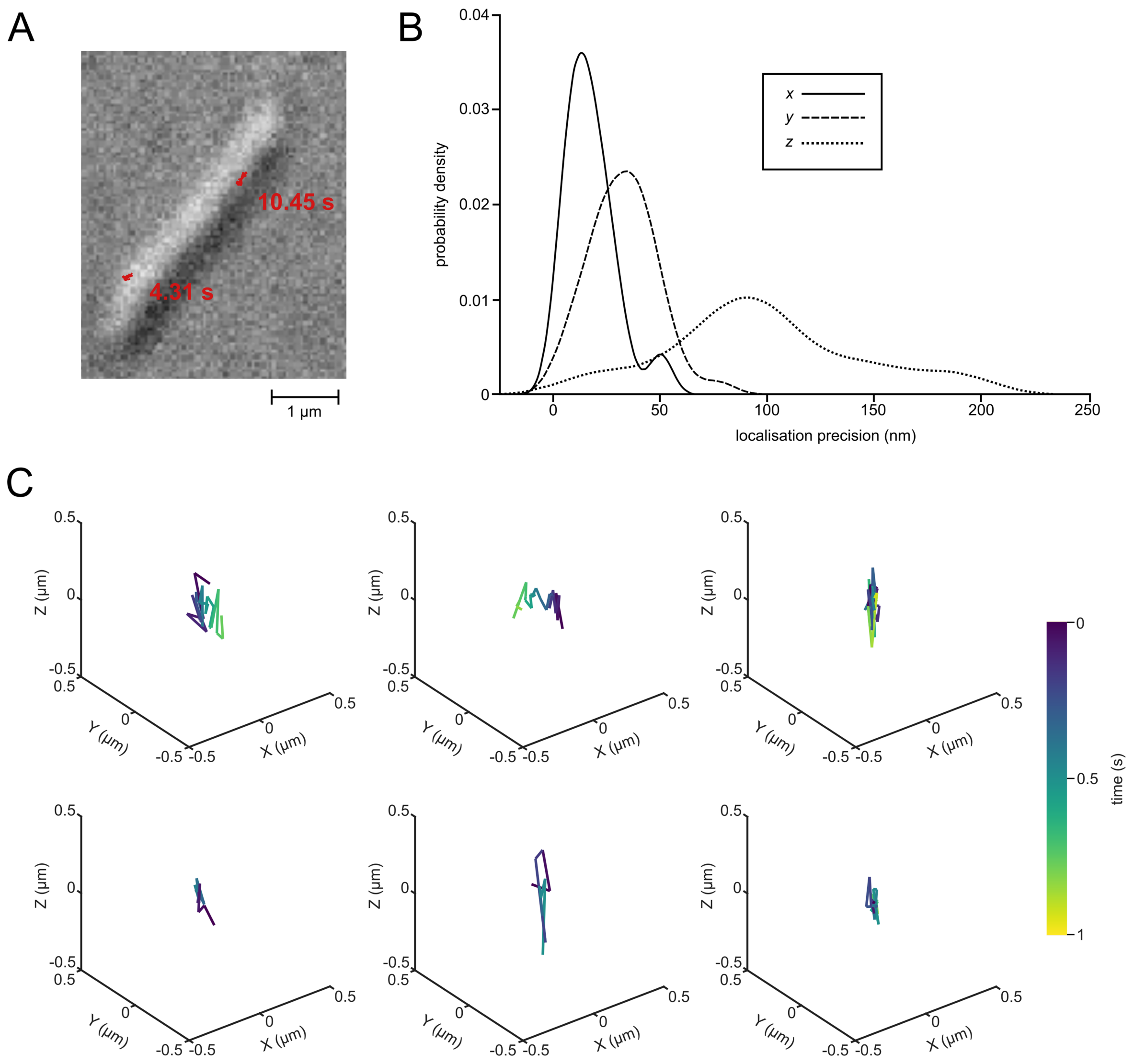


**Figure S5**

Imaging a HaloTag fusion of the stationary protein SprE, a component of the Type 9 Secretion System translocon which is stationary in the outer membrane (Lauber et al. 2018). (A) Tracks of SprE overlaid on brightfield images of a cell. Long stationary traces (acquired across 10.45 and 4.31 s respectively for upper and lower trajectories) rule out the possibility of cell rotation in the agar. (B) Probability density for localisation precision in each dimension calculated from SprE trajectories. (C) 6 examples of trajectories used for calculating localization precision. Uncertainties in *y* exceed those in *x* because parallax splitting of the back-focal-plane in the *y*-axis increases the *y*-axis width of the point-spread-functions. Uncertainties in *z* are greater still because they are calculated from a difference in the *y* localisations.

**Figure S6**

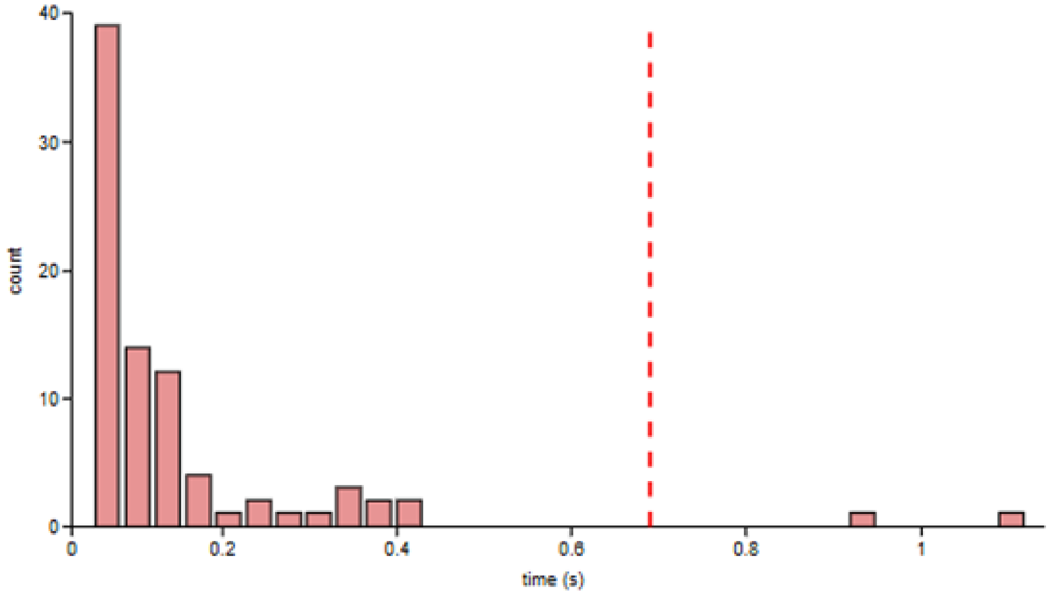


Control data tracking dye molecules non-specifically attached to cells: histogram of fluorescence trajectory durations in *ΔporV* mutant cells lacking the HaloTag ligand, incubated with Janelia Fluor 646 dye and following the same protocol as for labelled SprB. The vertical dashed red line marks the 20-frame threshold applied to SprB data to exclude non-specifically attached dye molecules. Only 2 of 83 trajectories (2.4%, in 47 cells from 3 cultures, both non-moving) exceeded the threshold, indicating the lifetime of non-specific attachment of dye molecules.

**Figure S7**

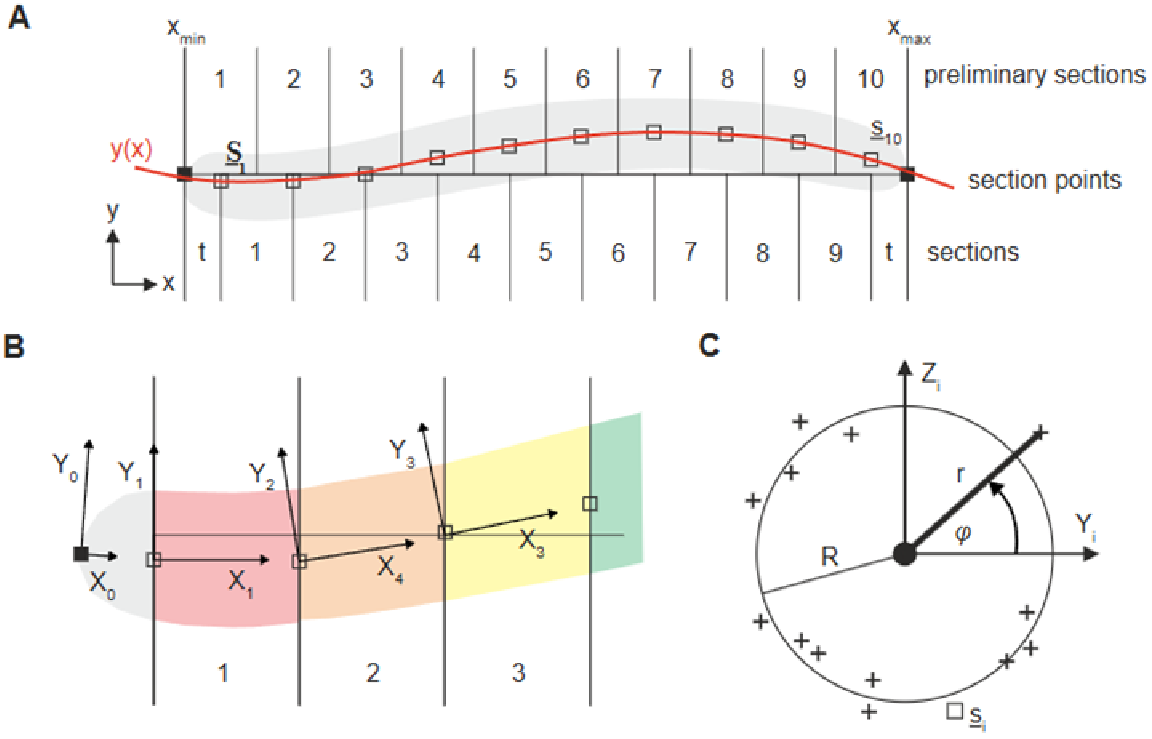


Illustration of the process of mapping 3D localizations to cylindrical co-ordinates for cell unrolling (Methods). A. Fitted cell mid-line (red), 10 preliminary sections, and 11 sections, corresponding to steps 1-4 in Methods. B) Local axes following the cell mid-line, and corresponding sections (coloured), corresponding to steps 5-6 in Methods. C) Fitting and transformation to local cylindrical co-ordinates, corresponding to steps 7,8 in Methods.
